# Overexpression of hypoxia-inducible factor 1 alpha improves immunomodulation by dental mesenchymal stem cells

**DOI:** 10.1101/141218

**Authors:** VG. Martínez, I. Ontoria-Oviedo, CP. Ricardo, SE. Harding, R. Sacedón, A. Varas, A. Zapata, P. Sepúlveda, A. Vicente

## Abstract

**Background:** Human dental mesenchymal stem cells (MSCs) are considered as highly accessible and attractive MSCs for use in regenerative medicine, yet some of their features are not as well characterized as in other MSCs. Hypoxia-preconditioning and hypoxia inducible factor 1 alpha (HIF-1 alpha) overexpression significantly improve MSC therapeutics, but the mechanisms involved are not fully understood. In the present study, we characterize immunomodulatory properties of dental MSCs and determine changes in their ability to modulate adaptive and innate immune populations after HIF-1 alpha overexpression.

**Methods:** Human dental MSCs were stably transduced with GFP (MSCs) or GFP-HIF-1 alpha lentivirus vectors (HIF-MSCs). Hypoxic-like metabolic profile was confirmed by mitochondrial and glycolysis stress test. Capacity of HIF-MSCs to modulate T cell activation, dendritic cell differentiation, monocyte migration and polarizations towards macrophages and NK cell lytic activity was assessed by a number of functional assays in co-cultures. Expression of relevant factors were determined by PCR analysis and ELISA.

**Results:** While HIF-1 alpha overexpression did not modify inhibition of T cell activation by MSCs, HIF-MSCs impaired dendritic cell differentiation more efficiently. HIF-MSCs induced also higher attraction of monocytes, which differentiate into suppressor macrophages, and exhibited enhanced resistance to NK cell-mediated lysis, which support the improved therapeutic capacity of HIF-MSCs. HIF-MSCs also displayed a pro-angiogenic profile characterized by increased expression of *CXCL12/SDF1* and *CCL5/RANTES* and complete loss of *CXCL10/IP10* transcription.

**Conclusions:** Immunomodulation and expression of trophic factors by dental MSCs make them perfect candidates for cell therapy. Overexpression of HIF-1 alpha enhances these features and increases their resistance to allogenic NK cell lysis and, hence, their potential *in vivo* lifespan. Our results further support the use of HIF-1 alpha-expressing dental MSCs for cell therapy in tissue injury and immune disorders.

## Background

Among different sources of MSCs, dental MSCs (dental pulp, periodontal ligament, apical papilla, dental follicle and gingival tissue) are currently considered as one of the most attractive MSCs for use in engineering tissue and regenerative medicine. Great advantages presented by dental MSCs include accessibility, easy *in vitro* expansion, low immunogenicity and ability to differentiate into mesoderm-derived tissues [1]. Low frequency of *in vivo* differentiation of transplanted MSCs in experimental models [2–5] indicates that MSCs promote tissue repair through paracrine modulation of surrounding cells. Cell populations normally targeted by MSCs include resident progenitor cells, endothelial cells and immune cells [6].

Many efforts are currently focused on furthering the therapeutic potential of MSCs. In this regard, we and others have previously demonstrated that hypoxia and, more specifically, overexpression of hypoxia-inducible factor-1 (HIF-1) by genetic engineering potentiates the therapeutic properties of MSCs [7]. Owing to the critical role that the immune response plays in most of the pathologies targeted by MSC therapy, a better understanding of the effects of hypoxia over the immunomodulatory capacity of MSCs is of great importance.

In the present study, we present a comprehensive characterization of the *in vitro* immunomodulatory properties and expression of immune and trophic factors in dental MSCs. Besides, we demonstrate that overexpression of HIF-1 alpha improves several of these features, enhancing also their resistance to NK cell-lytic activity.

## Methods

All procedures were approved by the Instituto de Salud Carlos III and institutional Ethical and Animal Care Committees.

### Vector production, lentiviral transduction and mesenchymal stem cells culture

MSCs from human dental pulp were expanded as described [8] and transduced with lentiviral vectors pWP-GFP (MSCs) or pWP-HIF-GFP (HIF-MSCs) as described [7]. Transduction efficiency was evaluated by flow cytometry (Coulter EPICS XL flow cytometer; Beckman Coulter) to determine the percentage of GFP-positive cells. Percentages of infection obtained were normally around 90%. Human dental pulp MSCs (n=3; Inbiobank) were cultured in DMEM low glucose (Sigma-Aldrich Spain) supplemented with 10% heat-inactivated fetal calf serum (FCS) (Gibco, Life Technologies, Thermo Fisher Scientific). Cells were grown until confluence and subsequently subcultured up to 12 passages. For stimulation assays, 5×10^4^ MSCs were seeded in 6-well flat-bottom culture plates and after 3 days, media were replenished and supplemented or not with 10 ng/ml of rhIFN-gamma (Invitrogen). After 15 hours, cells were lysed for subsequent analysis by quantitative-PCR (qPCR).

### Seahorse extracellular metabolic-flux assays

The Seahorse extracellular flux analyzer XFp (Seahorse Bioscience) was used to perform the Mitochondrial and Glycolysis Stress Tests. Detailed conditions in supplementary material.

Mitochondrial Stress Test: four basal oxygen consumption rate (OCR) measurements were taken. Subsequently, mitochondrial membrane modulators oligomycin (1 μM), followed by FCCP (1 μM) and lastly antimycin A/rotenone (both 1 μM) were injected (four measuring cycles each). Basal respiration, ATP production, maximal respiration, spare respiratory capacity and proton leak were defined by the differences between average measurements 4–16, 4–8, 12–16, 4–9 and 8–16, respectively. Non-mitochondrial respiration was defined by average measurement 16.

Glycolysis Stress Test: the extracellular acidification rate (ECAR) measurements were taken. Subsequently, glucose (10 mM), followed by oligomycin (1 μM) and 2-deoxy-glucose (2-DG, 100 mM) were injected (four measuring cycles each). Basal glycolysis, glycolytic capacity and glycolytic reserve were calculated by the differences between average measurements 4–8, 12–16, 8–12, respectively. Non-glycolytic acidification was defined by average measurement 16.

In both assays, total protein per well was quantified with the BCA Protein Assay Kit for normalization.

### Isolation and culture of human blood cells

Buffy coats of healthy donors were obtained after informed consent (Centro de Transfusión de la Comunidad de Madrid, Spain) and peripheral blood mononuclear cells (PBMCs) were isolated by density gradient centrifugation with Lymphocyte Isolation Solution (Rafer). Monocytes and NK cells were isolated by positive and negative magnetic separation (MiltenyiBiotec). Non-adherent T lymphocyte-enriched cell suspensions were obtained by nylon wool purification. Percentage of CD3^+^ cells was always above 85%.

CD14^+^ cells (5×10^5^) were cultured in 6-well flat-bottom culture plates for 6 days in 50% DMEM/10% FCS and 50% complete medium, consisting in RPMI (Lonza) supplemented with 10% FCS, 1 mM pyruvate, 2 mM glutamine, 100 U/ml penicillin and 100 μg/ml streptomycin (all from Sigma-Aldrich Spain), plus 20 ng/ml rhGM-CSF and 20 ng/ml rhIL-4 (Invitrogen) to generate immature monocyte-derived dendritic cells (MoDCs), or 5 ng/ml rhGM-CSF to generate monocyte-derived macrophages. In MoDC cultures, half of the medium was replaced by fresh medium and rhGM-CSF/rh IL-4 on days 2 and 4 of culture. In MDM cultures, additional 5 ng/ml rhGM-CSF was added every two days and 10 ng/ml of LPS (Invitrogen) was added for additional 24 hours of culture. When indicated, 5×10^4^ MSCs (MSC:Monocyte ratio 1:10) were seeded in 6-well flat-bottom culture plates, monocytes were added the day after and co-cultures were performed as stated above.

Freshly isolated NK cells were cultured in 24-well flat-bottom culture plates for 36 hours in 50% DMEM/10% FCS and 50% complete media supplemented with 10 ng/ml rhIL-12 and 10 ng/ml rhIL-15 (MiltenyiBiotec). As a negative control, NK cells were cultured in media alone. MSC-NK cell co-cultures were performed at MSC:NK cell ratios of 1:10 and 1:20, seeding MSCs the day before and keeping a number of 3×10^4^ MSCs.

For T cell activation, enriched T cell suspensions (3×10^5^) were cultured in 24-well flat-bottom culture plates for 4 days with immobilized anti-human CD3 (HIT3a) (10 μg/ml) and soluble anti-human CD28 (CD28.2) (1 μg/ml) monoclonal antibodies (BioLegend). As a negative control, T cell-enriched cells were cultured in media alone. MSC-T cell co-cultures were performed at a MSC:T cell-enriched ratio of 1:10, seeding MSCs the day before.

### Flow cytometry

See Supplementary Table 2 for a list of antibodies used. Immunofluorescence staining was performed as described [9]. For intracellular staining, brefeldin A (BioLegend) was added for the final 4 hours of culture and cells were treated with Fixation/Permeabilization Solution Kit (BD Biosciences). All flow cytometry analyses were conducted in a FACSCalibur flow cytometer (BD Biosciences) from the Centro de Citometría y Microscopía de Fluorescencia, Complutense University of Madrid.

### Proliferation assays

Before culture, T lymphocyte-enriched cell suspensions were labeled with 5μM CFSE (BioLegend). Proliferation of T cells (CD3^+^/CD4^+^ and CD3^+^/CD8^+^) was determined after 4 days by the CFSE dilution method. T cells were gated according to forward and side scatter.

### Apoptosis assays

The proportion of apoptotic MSCs was determined by incorporation of propidium iodide. MSC death was calculated as percentage of CD90^+^/propidium iodide^+^ cells.

### Migration assays

25×10^3^ MSCs were seeded in 24-well flat-bottom culture plates. The next day, culture media with 10 ng/ml rhIFN gamma was added and 6.5 mm insert, 8.0 μm polycarbonate membrane Transwell permeable supports (Corning Life Sciences) were mounted. 10^6^ PBMCs were added into inserts. Transwell cultures without MSCs in the lower chamber were used as controls. Cells were cultured for 14 hours and migrated cells (present at the lower chamber) were collected and stained for CD14. Number of CD14^+^ cells were counted on a FACSCalibur flow cytometer and gated according to forward/side scatter, and lack of CD90 expression. Monocyte migration (MM) was calculated as follows:

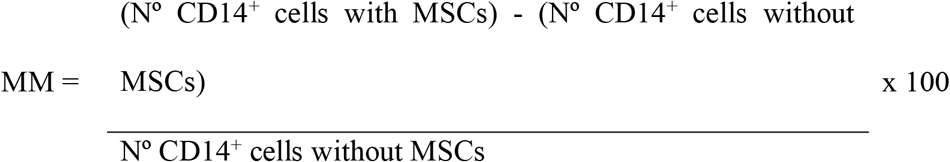

### PCR analysis

RNA isolation was performed using Absolutely RNA Microprep kit (Stratagene Cloning Systems, Agilent Technologies, Santa Clara, CA, USA), including DNase I digestion. Total cDNA was synthesized by High Capacity cDNA Reverse Transcription Kit (Applied Biosystems, Thermo Fisher Scientific). Taq-man assays (Applied Biosystems) employed for real-time PCR are in the supplementary material (Supplementary Table 2). All PCR reactions were set in duplicates using the TaqMan Gene Expression Master Mix (Applied Biosystems). Amplifications, detections, and analyses were performed in a 7.900HT Fast Real-time PCR System (Centro de Genómica, Complutense University, Madrid, Spain). The Delta CT method was used for normalization to GNB2L1 mRNA.

### Cytokine measurements

Levels of IFN gamma (R&D Systems), TNF-alpha and IL-10 in culture supernatants were assayed by ELISA (BioLegend). Production of CCL2/MCP-1 was measured by cytometric bead array (CBA, BD Biosciences).

### Statistical analysis

Student t test was used. Values of p≤0.05 (*/#) and p≤0.01 (**/##) were considered to be statistically significant.

## Results

### Efficient transduction of functionally active HIF-1 alpha in dental MSCs

MSCs from dental pulp were expanded and transduced with the lentiviral vectors pWP-GFP (MSCs) or pWP-HIF-GFP (HIF-MSCs). Efficient transduction of MSCs was confirmed by western blot and q-PCR, showing increased levels of HIF-1 alpha protein and mRNA in HIF-MSCs (figure 1A and 1B).

**Figure 1.**
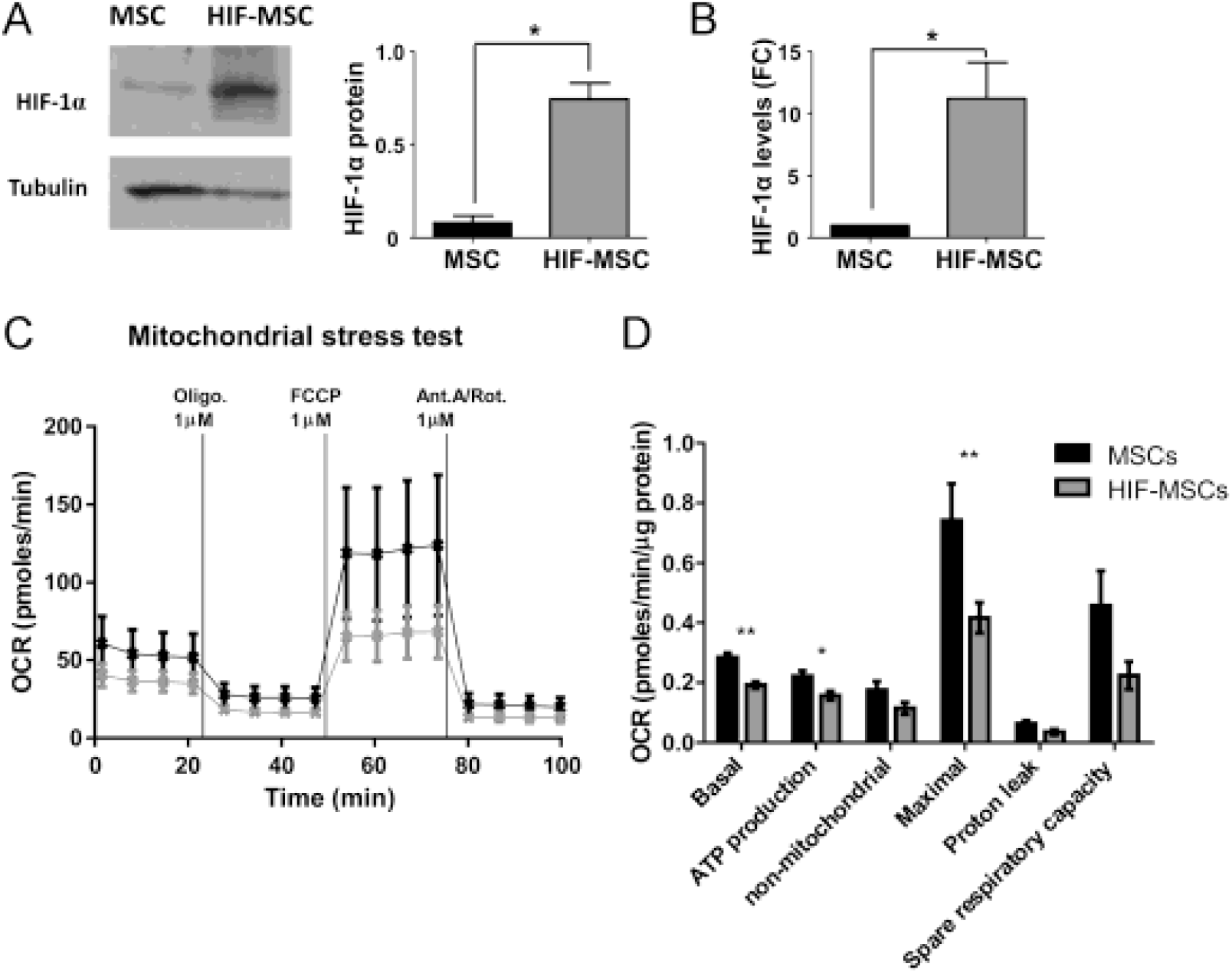
Overexpression of HIF-1 alpha and mitochondrial stress test in MSCs. A) Representative Western blot of HIF-1 alpha in MSCs and HIF-MSCs. Loading control was performed with Tubulin and levels of HIF-1 alpha protein expression were quantified by densitometry (right graph). B) Expression of the HIF-1 alpha mRNA determined by quantitative PCR. GNB2L1 was used as endogenous control. Date represent expression in HIF-MSCs relative to MSCs control (FC, fold change). Quantification of 4 independent experiments, data are represented as mean ± SE (*p≤0.05). C) Oxygen comsumption rate (OCR) response to Oligomycin (1 μM), FCCP (1 μM), and antimycin A/rotenone (both 1 μM) injections. HIF-MSCs (gray line) showed a lower OCR than MSCs (black line). D) Mitochondrial substrate utilization in HIF-MSCs (gray) and MSCs (black) measured in terms of basal respiration, ATP production, non-mitochondrial respiration, maximal respiration, proton leak and spare respiratory capacity. Data were normalized against the total amount of protein. Data are represented as mean ± SE of 6 independent experiments (*p≤0.05, **p≤0.01).

To investigate the impact of HIF-1 alpha overexpression in the metabolic state of MSCs, we performed Mitochondrial and Glycolysis Stress Tests to compare the OCR and glycolysis, respectively, between HIF-MSCs with control MSCs. HIF-MSCs presented lower basal respiration, ATP production and maximal respiration rates, as assessed by injections of 1μM Oligomicin, 1μM FCCP and 1μM of antimycin A/rotenone (figure 1C and 1D). Additionally, HIF-MSCs tended to show a higher ECAR upon injection of 10mM Glucose (Glycolysis) and 1μM Oligomycin (Glycolytic capacity), indicating a possible effect of HIF-1 alpha in enhancing glycolysis (Supplementary figure 1). Altogether, protein and mRNA levels and metabolic changes demonstrate that HIF-1 alpha is functionally overexpressed in dental HIF-MSCs.

### Effects of HIF-1 alpha overexpression in the modulation of the adaptive immune response by dental MSCs

T cells and DCs are main players in adaptive immunity, with T cells acting as main effectors when a deleterious inflammatory response is developed. In addition, impairment of T cell response is a well-established feature of MSCs [10,11]. As a first approach, we investigated the impact of HIF-1 alpha expression on the ability of MSCs to inhibit TCR-triggered activation of T cells. When T cells were activated in the presence of MSCs, proliferation of both CD4^+^ and CD8^+^ T cells was dramatically reduced, regardless HIF-1 alpha overexpression by MSCs (figure 2A and 2B). As expected, levels of IFN-gamma secreted by activated T cells were severely reduced in the presence of both MSCs and HIF-MSCs (figure 2C).

**Figure 2.**
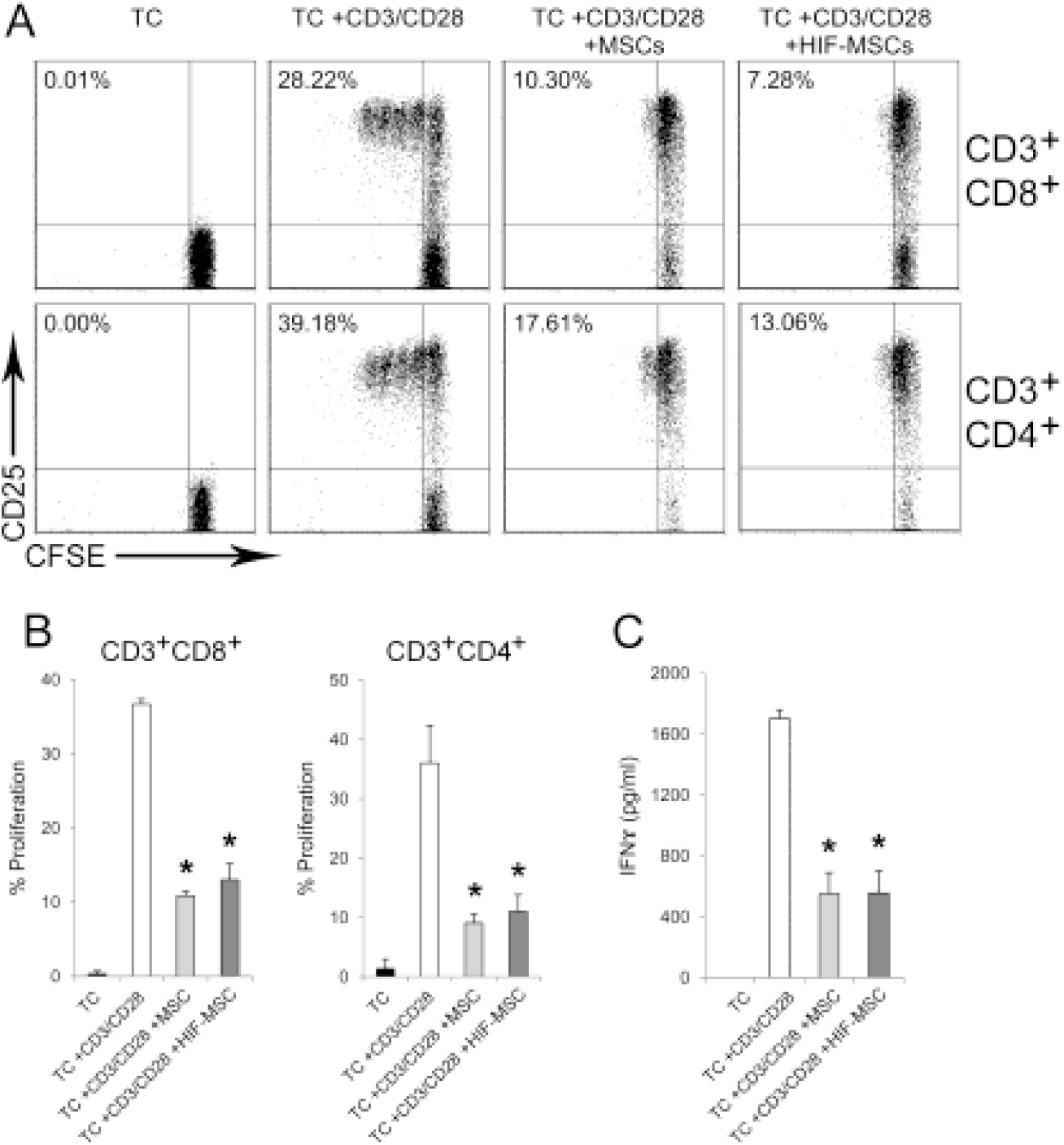
HIF-1 alpha overexpression does not modify the ability of MSCs to inhibit T cell activation. T cell-enriched peripheral blood cells were stained with CFSE and activated with anti-CD3/anti-CD28 monoclonal antibodies in the presence or absence of MSCs (light grey) and HIF-MSCs (dark grey). T cells were cultured in media alone as a negative control. After four days, expression of CD25 and proliferation of T cell subsets was determined by flow cytometry. A representative experiment (A) and the mean ± SD of three independent experiments (B) are shown. T cells were gated according to forward/side scatter characteristics and expression of CD3, CD4 and CD8. C) Supernatants of cultures were collected after four days and assayed for IFN gamma production. Data are presented as mean ± SD of three independent experiments. (*p≤0.05; MSCs and HIF-MSCs vs No MSCs).

We next tested whether HIF-1 alpha overexpression could improve the ability of MSCs to dampen DC differentiation. Dental MSCs negatively affected the generation of CD14^−^CD1a^+^ MoDCs from monocytes. Importantly, the inhibition of MoDC differentiation was significantly higher in HIF-MSC co-cultures, with a constant reduction in the percentage of CD14^−^CD1a^+^ MoDCs observed in control co-cultures (figure 3A and 3B).

**Figure 3.**
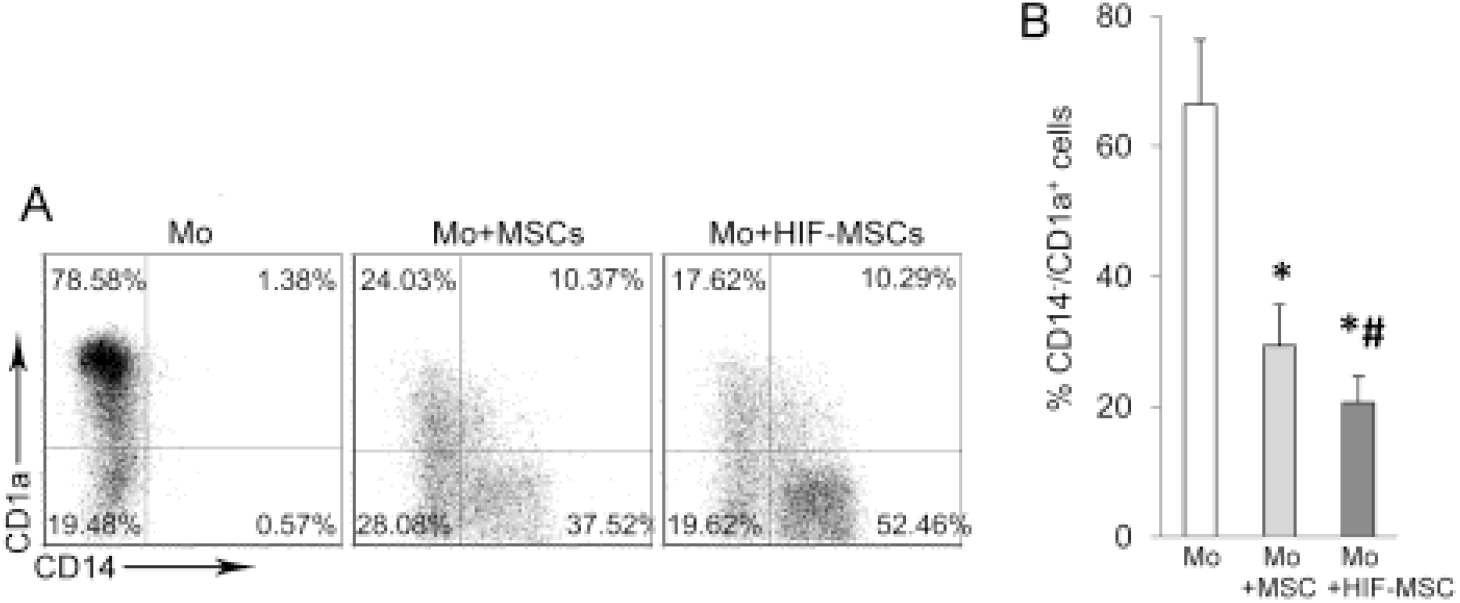
Increased capacity of HIF-1 alpha expressing MSCs to inhibit dendritic cell differentiation. Monocytes were differentiated towards dendritic cells with rhGM-CSF and rhIL-4 alone (Mo) or in the presence of MSCs (light grey) and HIF-MSCs (dark grey). After seven days, cells were harvested and expression of CD14 and CD1a was analyzed by flow cytometry. A representative experiment (A) and mean ± SD of four independent experiments (B) are shown. MSCs were excluded of the analysis by CD90 expression. (*p≤0.05; MSCs and HIF-MSCs vs No MSCs; #p≤0.05; MSCs vs HIF-MSCs).

### Overexpression of HIF-1 alpha by MSCs increases recruitment of monocytes which acquire immunosuppressive properties

Generation of pro-inflammatory macrophages from blood monocytes in response to environmental factors is a key phenomenon in the development of inflammation and tissue damage [12]. Our results show that the majority of monocytes cultured with both types of MSCs exhibited a CD14^high^/CD163^high^ phenotype (figure 4A and 4B). This phenotype has been previously associated to suppressor or M2 features in macrophages [13]. In this line, we found that the production of TNF-alpha was severely impaired by the presence of both MSCs and HIF-MSCs while IL-10 levels were remarkably increased in these co-cultures (figure 4C).

**Figure 4.**
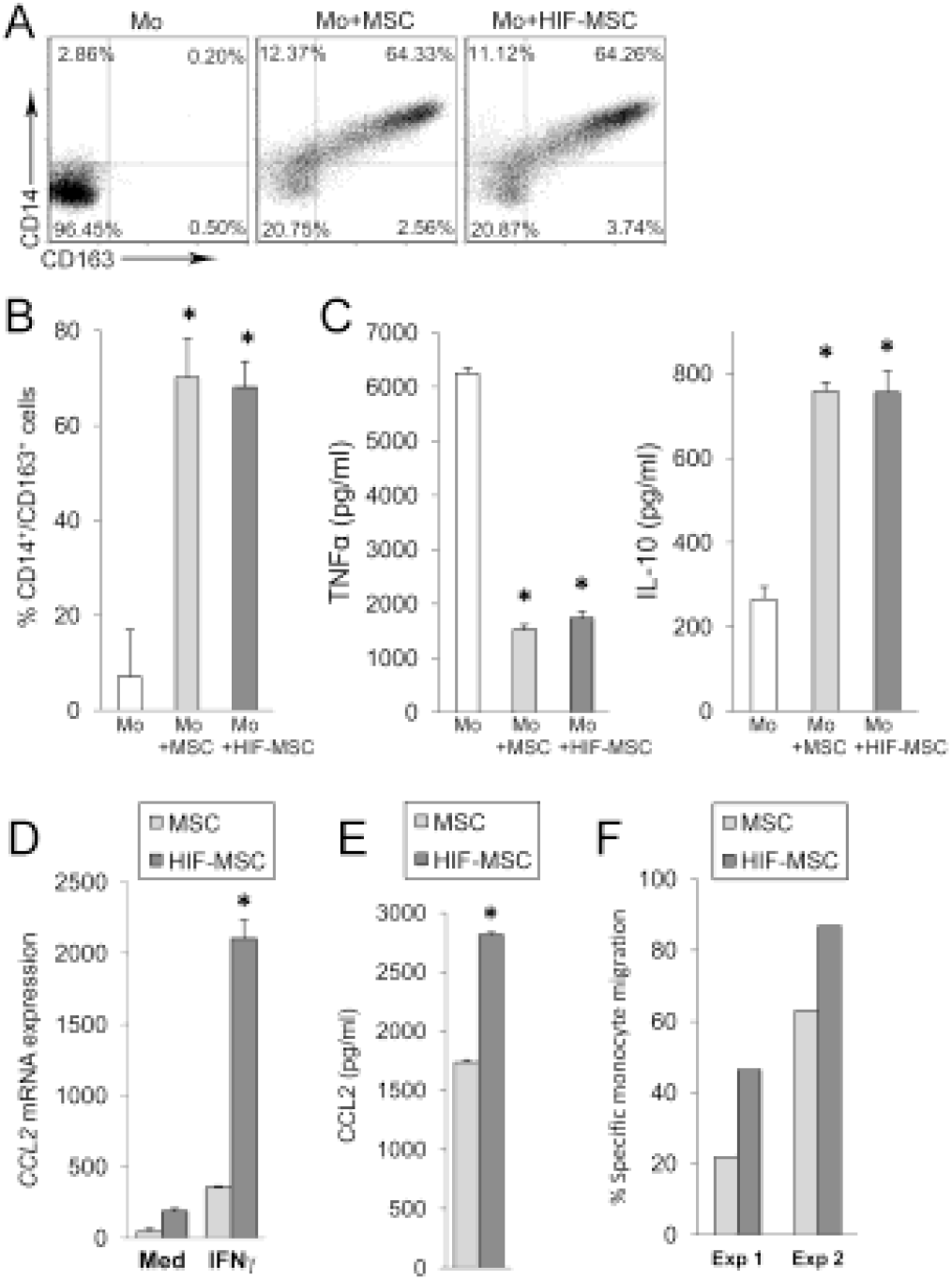
Monocytes acquire alternatively activated features in the presence of MSCs and are more efficiently attracted by HIF-MSCs. Monocytes alone (Mo) or in the presence of MSCs (light grey) and HIF-MSCs (dark grey) were cultured with rhGM-CSF to induce macrophage differentiation. After seven days, expression of CD14 and CD163 was determined by flow cytometry. A representative experiment (A) and mean ± SD of three independent experiments (B) are shown. MSCs were excluded from the analysis by CD90 expression. C) After 7 days of culture, LPS was added for 24 hours and supernatants were assayed for TNF alpha and IL-10 production. Data are presented as mean ± SD of three independent experiments. (*p≤0.05; MSCs and HIF-MSCs vs No MSCs). D and E) MSCs were cultured until 80% confluence, then fresh medium alone (Med) or supplemented with IFN gamma (IFNγ) was added. After 15 hour cells and supernatants were harvested. Transcript (D) and protein levels (E) for CCL2 were determined by quantitative PCR and CBA respectively. GNB2L1 was used as endogenous control. Mean ± SD of four samples pooled from two independent experiments is shown. (*p≤0.05; MSCs vs HIF-MSCs). F) Transwell (8 μm) cultures were performed as stated in the materials and methods section. After 14 hours, migrated cells (present at the lower chamber) were collected and stained for CD14. MSCs were excluded from the analysis by CD90 expression and monocytes were gated according to forward/side scatter characteristics and expression of CD14, and counted in a FACSCalibur flow cytometer.

These results suggested that the recruitment of monocytes to the site of inflammation could be advantageous in patients transplanted with dental MSCs. In this regard, CCL2/MCP-1 is the main chemokine driving monocyte extravasation. We found that HIF-MSCs exhibited augmented transcription of *CCL2/MCP-1* and that this difference was further enhanced in response to IFN-gamma (figure 4D). Coherently, secretion of CCL2/MCP-1 was significantly higher in IFN-gamma-stimulated HIF-MSCs (figure 4E), while basal levels of this chemokine were hardly detected (data not shown). Migration assays in the presence of IFN-gamma showed that the higher production of CCL2/MCP-1 observed in HIF-MSCs was accompanied by enhanced migration of monocytes (figure 4F).

### HIF-1 alpha overexpression confers resistance to NK cell-mediated lysis

Poor mid-term persistence of transplanted MSCs limits the therapeutic impact of MSCs, most notably in myocardial infarction cell therapy [14]. NK cells have been postulated as key mediators in this process, since they efficiently lyse autologous and allogeneic MSCs, participating in the clearance of these cells after transplantation, also accompanied by IFN gamma secretion [15]. Consistent with the literature, we found that degranulation of activated NK cells was increased in the presence of dental MSCs (figure 5A). Nevertheless, in HIF-MSC co-cultures NK cell degranulation was significantly impaired. In accordance, HIF-MSCs exhibited a marked resistance to NK cell-mediated death (figure 5B). Also importantly, increased IFN-gamma production by NK cells was strongly reduced by HIF-MSCs (figure 5C).

Interestingly, increased resistance of HIF-MSCs to NK cell-mediated lysis was not associated to differential modulation of the NK cell effector molecules granzyme B and perforin (figure 5D). Activation of the receptors Natural Killer Group 2D (NKG2D) and NKp30 has been shown to be critical for optimal lysis of MSCs by activated NK cells [15]. Nevertheless, we found no differences in the expression of NKG2D and NKp30 between NK cells activated in the presence of MSCs or HIF-MSCs (figure 5D). On the other hand, analysis of the expression by MSCs of the ligands for these NK cell activating receptor revealed a reduction in the basal expression of *ULBP1* and *B7H6* in HIF-MSCs (figure 5E). These dissimilarities were strengthened after stimulation with IFN-gamma, a pro-inflammatory factor highly produced by activated NK cells, again with a reduced expression of *ULBP1, B7H6* and *ULBP2* on HIF-MSCs (figure 5F).

**Figure 5.**
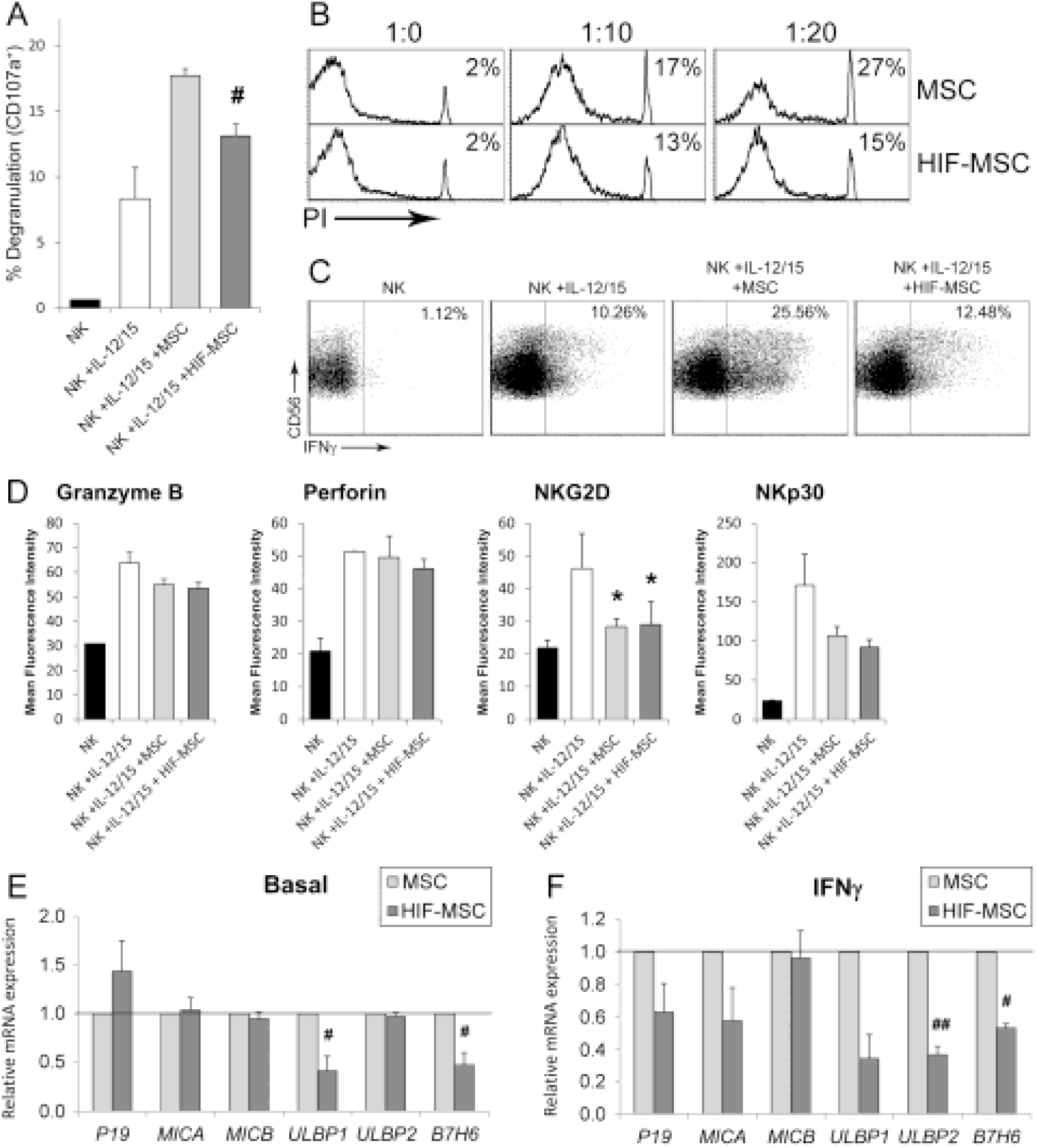
HIF-1 overexpression in MSCs confers resistance to NK cell-mediated lysis. NK cells were cultured in media alone (black bars) or with rhIL-12 and rhIL-15 in the absence (white bars) or presence of MSCs (light grey) and HIF-MSCs (dark grey). After 36 hours of culture, expression of CD107a (A), intracellular IFN gamma (C) and the indicated molecules (D) was analyzed in NK cells. MSCs were excluded from the analysis by CD90 expression. B) MSC-NK cell co-cultures were performed at the indicated MSC:NK cell ratios in presence of rhIL-12 and rhIL-15. After 36 hours late apoptosis of MSCs was determined by CD90 expression and propidium iodide incorporation. A representative experiment (B and C) and mean ± SD of three independent experiments (A and D) are shown. E and F) MSCs were cultured until 80% confluence was reached, then culture medium was replaced by fresh medium alone (E) or supplemented with IFN gamma (F). After 15 hours cells were harvested and expression of the indicated genes was determined by quantitative PCR. GNB2L1 was used as endogenous control. Date represent expression in HIF-MSCs relative to MSCs control. Mean ± SD of six to eight samples pooled from at least three independent experiments is shown. (*p≤0.05, MSCs and HIF-MSCs vs No MSCs; #p≤0.05, ##p≤0.005, MSCs vs HIF-MSCs).

### Enhanced immunosuppressive and pro-angiogenic features in HIF-1 alpha expressing MSCs

Following, we analyzed the expression of relevant factors regulating inflammation and tissue remodeling. Figure 6A represents the expression of immunomodulatory factors in control dental MSCs at basal levels and after IFN-gamma stimulation. As shown in the figure, IFN-gamma induced in MSCs the expression of *TLR3, TLR4, IL6* and, to a lesser extent, *COX2*. On the contrary, *LGALS1*, encoding for galectin 1, was greatly expressed by dental MSCs but not induced after IFN-gamma stimulation (figure 6A). Comparative analysis showed enhanced basal expression of the immunosuppressive molecules galectin 1 and IL-6 in HIF-MSCs (figure 6B, left panel). In an inflammatory context, mimicked by IFN-gamma stimulation, HIF-MSCs maintained a higher expression of *IL6* (figure 6B, right panel).

**Figure 6.**
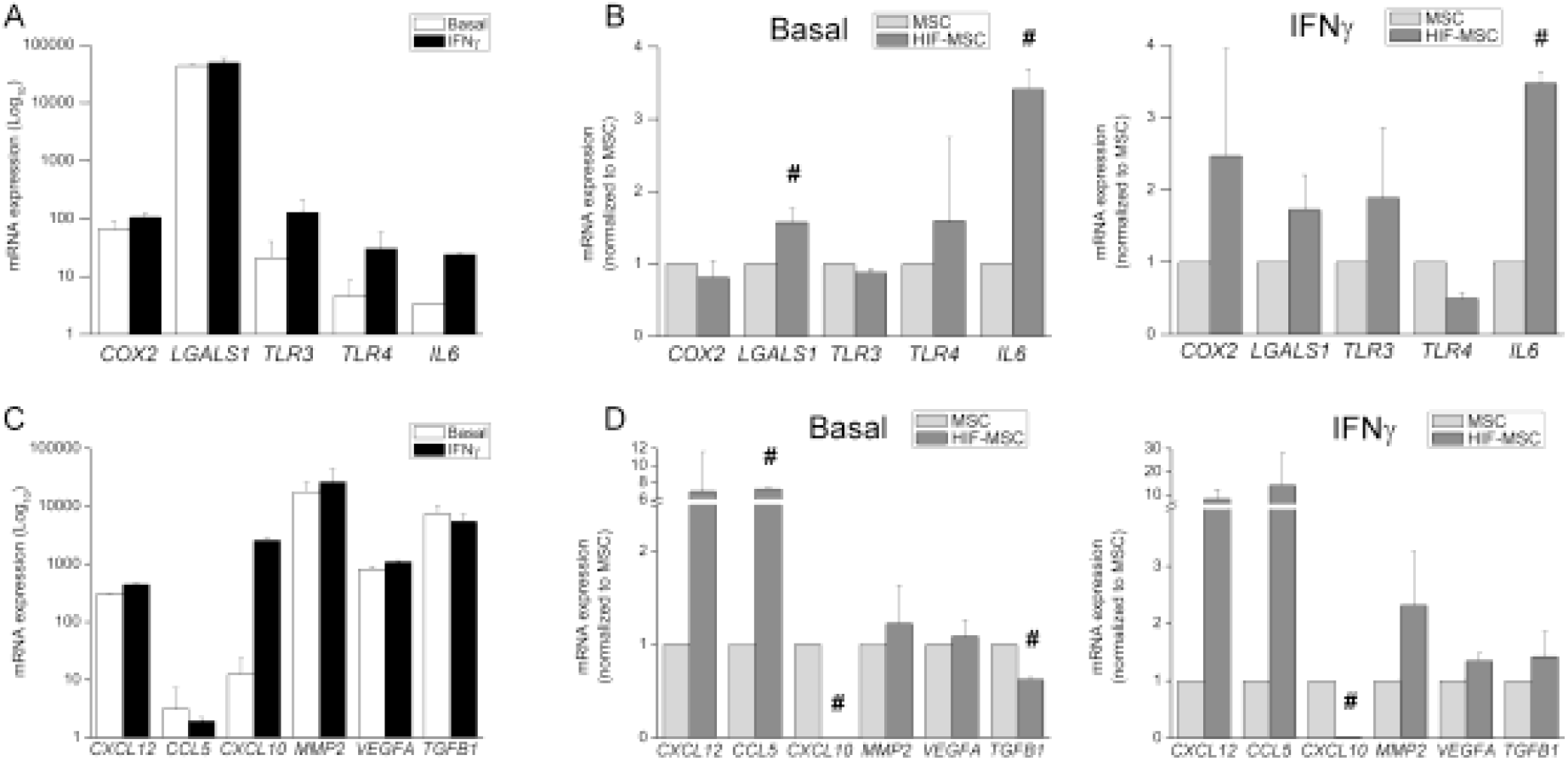
Immunomodulatory and pro-angiogenic profile induced by HIF-1 alpha expression in MSCs. MSCs were cultured until 80% confluence was reached, then culture medium was replaced by fresh medium alone (Basal) or supplemented with IFN gamma (IFNγ). After 15 hours cells were harvested and expression of the indicated genes was determined by quantitative PCR. GNB2L1 was used as endogenous control. A and C) Expression of the indicated genes in control MSCs. Note that logarithmic scale is used. B and D) Expression in HIF-MSCs relative to MSCs control. Mean ± SD of six to eight samples pooled from at least three independent experiments is shown. (#p≤0.05, MSCs vs HIF-MSCs).

We found also that dental MSCs differentially expressed a number of pro-angiogenic and trophic factors of which only *CXCL10* showed strong induction after IFN gamma treatment (figure 6C). Expression of *CXCL12* and *CCL5* was higher in HIF-MSCs both at basal levels and after addition of IFN-gamma, while the basal transcription of the pro-fibrotic factor *TGFB1* was reduced (figure 6D, left and right panels). Remarkably, overexpression of HIF-1 alpha caused a complete loss of *CXCL10* transcription in HIF-MSCs under basal and inflammatory conditions.

## Discussion

Dental MSC-based cell therapy is currently under extensive research in a wide range of pathologies in which inflammation plays a key role. These include wound healing, spinal cord injury and periodontitis [2,16,17]. While suppressor activities through T cell and macrophage responses have been previously described [18], the role of dental MSCs cells in multiple immune interactions remains elusive.

Our results show that inhibition of CD4^+^ and CD8^+^ T cell responses to TCR stimulation by dental MSCs is independent of HIF-1 alpha expression. This might be correlated with the lack of differences in the expression of COX2/PGE2, IL-10, and IDO (figure 5 and data not shown), main factors driving T cell inhibition by dental MSCs [19,20]. Recently, it was demonstrated that inhibition of proliferation of PBMCs by gingiva MSCs is increased after hypoxia preconditioning, with the participation of IL-10 and FasL [21]. Since total PBMCs were used in this experiments, it is difficult to know whether T cells were directly targeted by hypoxic gingiva MSCs. Accordingly, inhibition of T cell proliferation by adipose tissue-derived MSCs is not altered under hypoxic conditions [22].

On the other hand, we found that HIF-1 alpha overexpression induces higher expression *IL6*, a known negative regulator of DC differentiation and function [23,24]. Enhanced expression of *IL6* has also been shown in hypoxia-conditioned MSCs from bone marrow [25]. This could explain that impairment of DC differentiation by dental MSCs is more efficient when MSCs express HIF-1 alpha.

We also discovered that HIF-MSCs produce higher levels of the chemokine *CCL2/MCP1* under inflammatory conditions and promote greater attraction of monocytes. This is in agreement with recent data showing that conditioned medium from hypoxic bone marrow-derived MSCs enhances monocyte migration and recruitment of F4/80^+^ macrophages during wound healing [25]. Recent reports show that dental MSCs induces suppressor macrophage differentiation with the participation of different factors including IL-6 and CCL2/MCP-1 [17,26]. Furthermore, we present here that monocytes cultured with dental MSCs exhibit high IL-10 and low TNF-alpha production after LPS/TLR-4 stimulation. It is important to note that TLR-4 agonists are normally present after tissue damage [27]. Owing to the beneficial role normally attributed to anti-inflammatory macrophages [28], we propose that expression of HIF-1 alpha by MSCs would be advantageous for the resolution of inflammation through enhanced recruitment and differentiation of suppressor macrophages.

NK cell-mediated lysis of MSCs might be linked to the poor mid-term persistence of transplanted MSCs observed *in vivo* [14,15,29]. Here, we confirm these observations showing that NK cells are capable of killing dental MSCs, along with increased production of IFN gamma. Nevertheless, HIF-MSCs exhibit increased resistance to NK cell-mediated lysis as shown by reduced MSC cell death, NK cell degranulation and IFN gamma production. Efficient NK cell-mediated lysis requires binding of the activating receptors expressed on NK cells, to their natural ligands, expressed in MSCs [15]. Consistently, we show that HIF-MSCs exhibit a significantly reduced expression of the NK cell activating receptor ligands ULBP1 and B7H6 [30]. Interestingly, this mechanism is frequently exploited by tumor cells [31–33]. These results are coherent with the increased engraftment and survival of hypoxia-conditioned MSCs after transplantation [34,35]. Considering that poor mid-term persistence of transplanted MSCs is a limiting factor for the therapeutic potential of MSCs [14], we provide here evidences which support that HIF-1 alpha expression could be a useful approach to overcome this limitation.

MSCs may exert tissue repair and regeneration through paracrine factors, acting on surrounding cells and directing cytoprotection and neovascularization [6]. We have found that HIF-MSCs exhibit enhanced expression of *CXCL12/SDF1* and *CCL5/RANTES*, factors widely known by their pro-angiogenic actions [36,37]. Strikingly, our results demonstrate that HIF-1 alpha expression in MSCs completely abrogates *CXCL10/IP10* expression. This factor can limit new vessel growth by inhibiting endothelial cell migration, and can induce involution of new vessels [38]. Similarly, enhanced angiogenic potential of hypoxia-conditioned MSCs has been shown in two different experimental models for ischemia [34,39].

## Conclusions

Hypoxia-preconditioning of MSCs significantly improves the therapeutics of MSCs. Furthermore, this procedure has been suggested for standardized protocols of MSC preparation for clinical applications [40]. The present work demonstrates that HIF-1 alpha overexpression confers to MSCs enhanced paracrine capacities which might participate in the previously described benefits of HIF-MSC therapy after myocardial infarction [7]. Furthermore, HIF-MSCs exhibit increased resistance to NK cell-mediated lysis, which will help to prolong their lifespan in the host. We believe that dental MSCs and genetic engineering to express HIF-1 alpha are powerful tools for furthering the therapeutic potential of cell therapy.

## Declarations

### Ethics approval and consent to participate

All procedures were approved by the Instituto de Salud Carlos III and institutional Ethical and Animal Care Committees. Buffy coats of healthy donors were obtained after informed consent (Centro de Transfusión de la Comunidad de Madrid, Spain).

### Consent for publication

Not applicable.

### Availability of data and material

All data generated or analysed during this study are included in this published article and its supplementary information files.

### Authors′ contributions

MVG, SR, VaA, ZA and ViA were responsible for design, performance and analysis/interpretation of the in vitro experiments. OI and SP were responsible for generation and analysis of MSC cell lines. RCP and HSE were responsible for metabolic analysis of MSCs. MVG wrote the manuscript. All authors read and approved the manuscript.

## Acknowledgements

Not applicable.

## Additional file 1: Supplementary Material

Word file (.docx) including:

1. Detailed methods: lentiviral production and transduction, metabolic assays and list of reagents for flow cytometry and PCR.
2. Supplementary figure 1: Results from glycolytic activity of MSCs.

